# ConfluentFUCCI for fully-automated analysis of cell-cycle progression in a highly dense collective of migrating cells

**DOI:** 10.1101/2023.10.20.563216

**Authors:** Leo Goldstien, Yael Lavi, Lior Atia

## Abstract

Understanding mechanisms underlying various physiological and pathological processes requires accurate and fully automated analysis of dense cell populations that collectively migrate, and specifically, relations between biophysical features and cell cycle progression aspects. A seminal tool that led to a leap in real-time study of cell cycle is the fluorescent ubiquitination-based cell cycle indicator (FUCCI). Here, we introduce ConfluentFUCCI, an open-source graphical user interface-based framework designed for fully automated analysis of cell cycle progression, cellular dynamics, and cellular morphology, in highly dense migrating cell collectives. Leveraging state-of-the-art tools, some incorporate deep learning, ConfluentFUCCI offers accurate nuclear segmentation and tracking using FUCCI tags, enabling comprehensive investigation of cell cycle progression at both the tissue and single-cell levels. We compare ConfluentFUCCI to the most recent relevant tool, showcasing its accuracy and efficiency in handling large datasets. Furthermore, we demonstrate the ability of ConfluentFUCCI to monitor cell cycle transitions, dynamics, and morphology within densely packed epithelial cell populations, enabling insights into mechanotransductive regulation of cell cycle progression. The presented tool provides a robust approach for investigating cell cycle-related phenomena in complex biological systems, offering potential applications in cancer research and other fields.

## Introduction

The cell cycle is a highly regulated process that governs the growth and division of cells. It consists of a series of stages, including DNA replication, chromosome segregation, and cytokinesis, which together result in the formation of two daughter cells. Disruptions in this process can lead to various physiological disorders, including cardiovascular disease, infection, inflammation, and cancer [1]. The progression of the cell cycle is tightly controlled by signaling pathways that are intracellular, integrated with mechanical cues that are intercellular. Such mechanical cues stem from the extracellular matrix and neighboring cells [2]. These mechanotransductive processes are likely to be highly pronounced when neighboring cells in a dense tissue apply substantial forces between one another during collective cellular migration [3]. In the context of collective migration, specifically of epithelial cells, the regulation of cell cycle progression is crucial for ensuring the coordinated movement of cells. Therefore, understanding the relationship between cell cycle progression and collective cellular migration is essential for elucidating the mechanisms that govern various pathophysiologies and developing new therapeutic strategies to treat diseases, involving abnormal cell migration.

Several studies have made significant advancements in the study of cell cycle progression during collective migration using the fluorescence ubiquitination-based cell cycle indicator (FUCCI) [4, 5]. FUCCI is a seminal tool that led to a phenomenal leap in real-time study of cell cycle progression under various organisms and protocols [6, 7].

The FUCCI sensor utilizes two distinct fluorescent proteins conjugated to specific regulating proteins at different cell cycle stages: MKO2 to cdt1 (G0/G1) and mAG1 to geminin (S/G2/M). As cdt1 decrease and geminin increase, the overlay of MKO2 and mAG1 channels display the nuclei in yellow (G1 to S transition), which marks cell cycle progression (Fig. 1). However, the inherent design of the FUCCI sensor, which indicates decreases or increases in cdt1 and geminin, makes both MKO2 and mAG1 fluorescent images very heterogeneous. These issues create significant computational challenges for image analysis. First, typical time-lapse imaging of collective migration experiments generates hundreds of images per position per sample. Coupled with high resolution cameras, modern imaging solutions often generate dozens to hundreds of gigabytes per experiment. At this scale we must make sure our computational approach is both appropriate and feasible. Of specific issue would be approaches that scale poorly with the number of detected nuclei or those that do not take advantage of multi-core processors or GPUs. Second, the FUCCI fluorescent signals exhibits a wide dynamic range both in space and time. This feature of the system makes selecting analysis parameters (such as thresholds, SNR bounds, etc.) challenging and error prone. Finally, the extreme variability and heterogeneity makes marking the transition from G1 to S tied to a specific analyst opinion, and thus non-robust.

**Figure 1.**
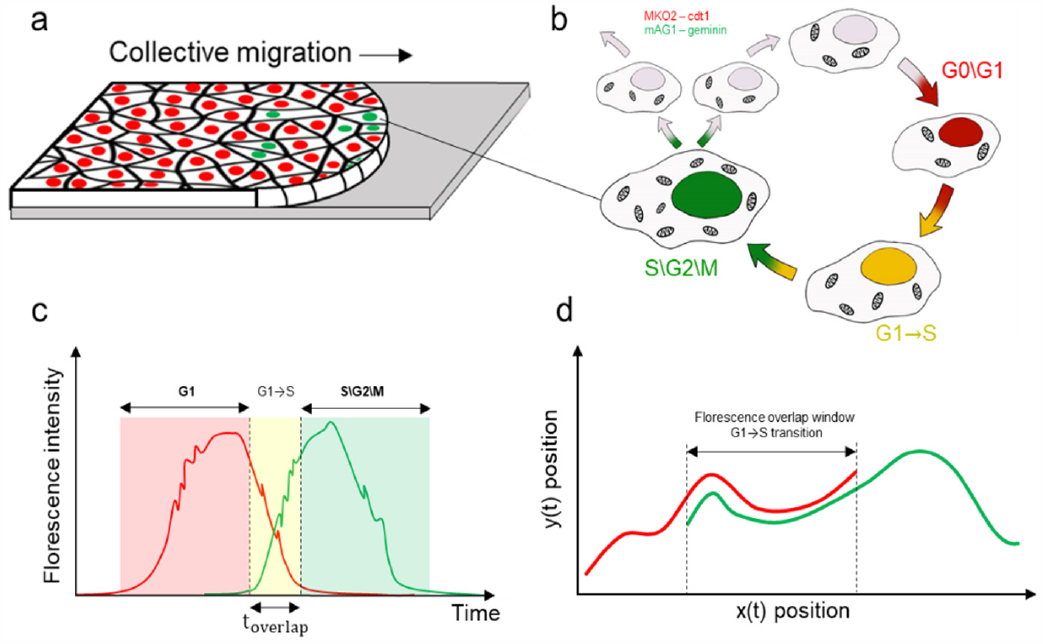
FUCCI monitoring during collective cellular migration. **a**, collective migration of confluent epithelial layer with cells tagged for FUCCI which, in **b** and **c** display different fluorescent colors as each cell progresses through the cell cycle. Newly formed cells are colorless. As G0\G1 progresses, the amount of cdt1 in the nucleus increase, and with it, the red fluorophore. When transitioning from G1 to S: 1) cdt1 concentration decreases, and with it, the red fluorophore gradually drops; 2) geminin concentration increases as shown by the rise in the green signal; 3) both fluorophores co-fluorecess and the cell appears yellow. When S progresses the red fluorophore disappears, while the appearance of green fluorophore continues to increase. Upon division, the green color is lost, with the disappearance of geminin. **d**. As each cell, among thousands in the layer, migrate, we track its position within each of the fluorescent channels, and while the two tracks are not perfectly aligned,

For image analysis of FUCCI time-lapse datasets, several semi-automatic analysis tools exist in which the user is involved in a laborious procedure of either manually annotating, or aiding the machine by meticulously supervising a prediction for the cell location and cell cycle status [8-10]. Out of these, however, even the most recently developed CPU-based tool (FUCCItrack, [8]) is inadequate, as we will show later, to handle in an accurate and timely fashion the analysis of huge amount of data in a typical experiment of collective epithelial migration within a dense tissue. Such a tissue is composed of thousands of extremely dynamic cells in each field of view (FOV). Hence, with multiple FOVs per tissue, in a multiwell culture plate, the analysis calls for an approach that is both accurate and scales to analyzing hundreds of gigabytes in a timely fashion. In this case, a Graphics Processing Unit (GPU) acceleration is essential for performing efficient image analysis that combines biological cell segmentation followed by position tracking. The main benefit for this is that many image analysis tasks often scale linearly on a GPU, facilitating the processing of massive datasets in a timely manner. GPU acceleration enables the execution of these tasks by leveraging thousands of cores to perform computations simultaneously, which is much faster than the sequential processing done by traditional CPU-based systems. Moreover, now days, GPU cloud servers provide accessible hardware for leveraging modern deep learning-based image analysis techniques, such as Convolutional Neural Networks (CNNs), which have shown great success in solving complex segmentation and tracking problems [11, 12].

Here we present ConfluentFUCCI, an open-source graphical user interface (GUI)-based framework. ConfluentFUCCI makes use of state-of-the-art Deep Learning algorithms to achieve accurate, fast and fully-automated analysis of cellular morphology, dynamics, and cell-cycle progression, for each FUCCI stained cell within thousands of collectively migrating cells. Our software uses an application programming interface (API) to match with Cellpose [13], an efficient Deep-Learning (DL) based segmentation tool with GPU acceleration, which facilitates accurate and flexible nuclear segmentation. Furthermore, embedded within ConfluentFUCCI is our new algorithm that: 1) interface with, and utilize, a dynamic nuclei tracking module in TrackMate [14] that perform nuclei tracking in the two separate FUCCI fluorescent channels (Fig. 1); 2) integrates tracking data from both channels, and assigns to each cell one continues track with compatible time dependent data that include cell and nucleus morphology, cell coordinates and their dynamics, and cell cycle state throughout the entire cell cycle. As such, ConfluentFUCCI leverages and integrates between two state-of-the-art tools that simplify the analysis of cell cycle data in migration assays, and defines a robust, and user-independent, criterion for G1 to S transition.

We will present a comparison with the most recently developed tool for such relevant analysis, and show that our computational scheme is more accurate than current solutions, applies to both sparse or dense cellular assays, and scales well to real-world conditions. We further show how we independently replicate the previously published data [4] of area-dependent cell-cycle transition in expanding tissue cultures of Madin-Darby canine kidney (MDCK) cells of epithelial origin. In addition, the approach proposed here provides a reproducible technique for tracking any live nuclei stain in dense epithelial populations using state-of-the-art tools, while providing specific benefits for analyzing FUCCI stained cultures.

## Materials and Methods

### The common route that we avoid

Until recently, the image analysis pipeline for biological cell shape and dynamic analysis typically involved a series of manual and semi-automated steps. The process began with acquiring images of cells using a microscope, followed by image preprocessing to remove noise and enhance contrast. Techniques such as Gaussian blurring and median filtering were commonly used to reduce noise and improve the quality of the images. Then, to separate individual cells from the background the three main classes of techniques that were employed included thresholding, edge detection, and region growing. Thresholding involves setting a threshold value to separate the foreground from the background in an image. Edge detection involves identifying the boundaries of cells by detecting changes in intensity between adjacent pixels. Region growing involves selecting a seed pixel and iteratively adding adjacent pixels that meet certain criteria to form a segmented region. Combined with these techniques, known morphological operations, such as erosion, dilation, and opening, were commonly employed. These were used to remove small objects or fill gaps in segmented regions, and to produce a finalized segmented image. To track a specific cell, or a segmented region, from one time point to another, a variety of tracking algorithms were used. These include algorithms such as nearest-neighbor, and Kalman filter, that relied on feature matching between frames to establish cell identity and its dynamics.

It is important to emphasizes that most mentioned techniques were often time-consuming, labor-intensive, prone to human errors, and did not scale well with large datasets. Hence, it was in the hands of the experienced bioimage analysts to creatively tailor from all mentioned techniques an image analysis pipeline that in most cases was suitable for a specific set of images, taken in a specific experiment, by a specific lab. Since we aspire to provide a robust tool to analyze the FUCCI data in a variety of flat culture conditions, the above approach that was commonly taken before [8, 10], does not suffice.

### The presented approach for FUCCI signal analysis in dense and dynamic tissues

In the case of multicellular migration, with thousands of cells per FOV, the FUCCI signal and its analysis raise multiple challenges. First, the nuclei are often clustered with potential overlapping in the image which frequently necessitates additional processing. Second, slightly out-of-focus regions in the image results in blurry nuclei boundary and difficulty in segmentation. Third, both MKO2 and mAG1 channels of the FUCCI marker display fluctuating intensities in both space and time. Spatial fluctuations result from fluorophore inhomogeneity within a nucleus, cell to cell shape variability, and the extremely close proximity of the nuclei in collective migration experiments. All these features make the common threshold selection, may it be global or local, very challenging for segmentation of all nuclei in the FOV in each time point. Furthermore, temporal fluctuations in a dense FUCCI system brings two main challenges. A problematic continuous tracking of any nucleus in either MKO2 or the mAG1 channel, and the ability to properly track the same nuclei simultaneously in both channels. Doing so allows us to define the exact overlap window between the MKO2 and mAG1 illuminations, and thus gives the ability to declare the transition from G1 to S in a user independent manner. That is, for example, one can define the entire overlap window (yellow phase), or perhaps its mid time point, to be the G1 to S time of transition. That way, the conventional approach of declaring the G1 to S transition point based on specific fluorescent intensity value, and when the nuclei appears to the user as yellowish [4, 8], is avoided.

### Implementation and integration approach

A typical problem with an available code or algorithm for bioimage analysis in the literature are malfunctions that arise when the tool is used on different computational environments then it was originally designed in. These include different machines, operating systems, auxiliary folders and so on. For this purpose, we wrapped ConfluentFUCCI in what is know as container. Containers facilitate packaging of an entire runtime environment, including the specific versions of libraries, configuration files, and datasets [15]. Additionally, containers provide a stable platform protecting against potential numerical instability when running in vastly different computational environments. Finally, the adoption of containerization of scientific software provides a path towards scalability and compatibility with cloud computing platforms, thus allowing access to extensive computational resources, which are often required for analyzing contemporary bioimage datasets. Thus, confluentFUCCI can be run on any operating system, with any type of available GPU or CPU. Yet, the biggest advantage we found in using a container was the ability to collect and integrate available state-of-the-art technologies such as deep learning, distributed computing, and modern webapps. Of specific interest to this work were NumPy (matrix operations) [16], Pandas (tabular data) [17], Dask (distributed computation) [18], Holoviews (graphs & figures) [19], Panel (WebApps) [20], Napari (multi-dimensinal image viewer) [21], TrackMate (object tracking) [14], and CellPose (machine learning based segmentation) [13]. CellPose and TrackMate are core tools used within confluentFUCCI.

CellPose developers trained a neural network model (termed U-net) on a diverse set of cell/nuceli images to create a relatively generic approach applicable to a diverse set of imaging modalities and cell types. This is what commonly referred to as Deep Learning (DL), which requires huge amount of computation tasks that are best handled with a GPU. CellPose has a native support of executing on GPUs which can significantly speed up segmentation. We used an available neural network trained model (cyto2) for nuclei detection in Cellpose and further improved it by training it on time lapse data with hundreds of MDCK-FUCCI cells imaged with a Zeiss microscope. The same model was able to detect both FUCCI marked cells for different cell lines, and for different imaging systems.

TrackMate is a mature and performant cell tracking software, which is distributed as a FIJI plugin. While both tools are open-source, interfacing between Python and Java is neither straightforward nor error free. Furthermore, TrackMate is GUI driven and difficult to operate programmatically. We overcame this by packaging both Fiji and TrackMate, as an easy to use container. We designed and included in the container a custom-built middleware for loading an image stack, setting up tracking settings and saving the results. ConfluentFUCCI uses TrackMate to track all detected nuclei in both fluorescent channels simultaneously. It then calculates a similarity metric to identify which tracks (or part of tracks) overlap in space & time in both green and red channels. This lets us definitively merge individual red/green tracks into one continuous track throughout the cell cycle. For each such track all the available information is saved. This include dynamics, area, shape, and cell cycle state.

### Workflow description and overview of functionalities

Fig. 2 describes the entire workflow of ConfluentFucci that is carried in a fully automatic manner from start to finish. The process begins with the two image stacks, red (MKO2) and green (mAG1), and the (neural network) trained model (based in cyto2 in CellPose) and its settings that are inputted for segmentation. The segmented stacks are individually passed to TrackMate (plugin in FIJI), together with the specific required settings. Segmented nuclei are tracked to identify red tracks and green tracks and are outputted as two XML files (also available in the user library). Thus far, every stack (red\green) was analyzed by itself. The program now searches which tracks in the red channel and the green cannel need to be paired and be stored as one continues track of one specific cell. This is made in the following two main programmatic steps. First, to alleviate calculation effort the code filters, from all possible pairs of green and red tracks, the pairs that are more probable to eventually be identified as belonging to the same cell. To do so, the image is divided to a grid of n*x*n equal squares, and only a cell-R_i_ from the red tracks, and a cell-G_j_ from the green tracks, that passed through the same square in the grid, at any time throughout their trajectories, are considered a probable pair. Second, for each probable pair R_i_-G_j_ we calculate a similarity metric S_ij_, defined as the average Euclidian distance between the centers of R_i_ and G_j_ nuclei throughout their trajectories. A pair R_i_-G_j_ for which S_ij_ is smaller than 2 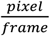is defined as belonging to the same cell. We find that the pre-filtering process in the two described steps works optimally for n=10. The program now takes an overlay of the green and red channels, and based on all nuclei centers preforms a Voronoi tessellation in order to approximate the area of each cell. The validity of this area approximation approach was previously demonstrated and shown to be accurate only in confluent and dense conditions [4, 22, 23], and can also be seen in Fig. 5a. Furthermore, not only that by definition each nuclei has a defined velocity vector after all steps above, the cellular velocity field for each position in the image is then also approximated. This is done by dividing the image to small bins, and calculate the average nuclei velocity in each bin. At the end of the process the user gets a table with all identified cells, their united trajectories, velocities, area, and the state in the cell cycle. Taken together, the described capabilities give ConfluentFUCCI users a unique approach to examine the relationship in space or time between, any desired morphological and dynamic aspects of the cell and its nucleus, and cell cycle status.

**Figure 2.**
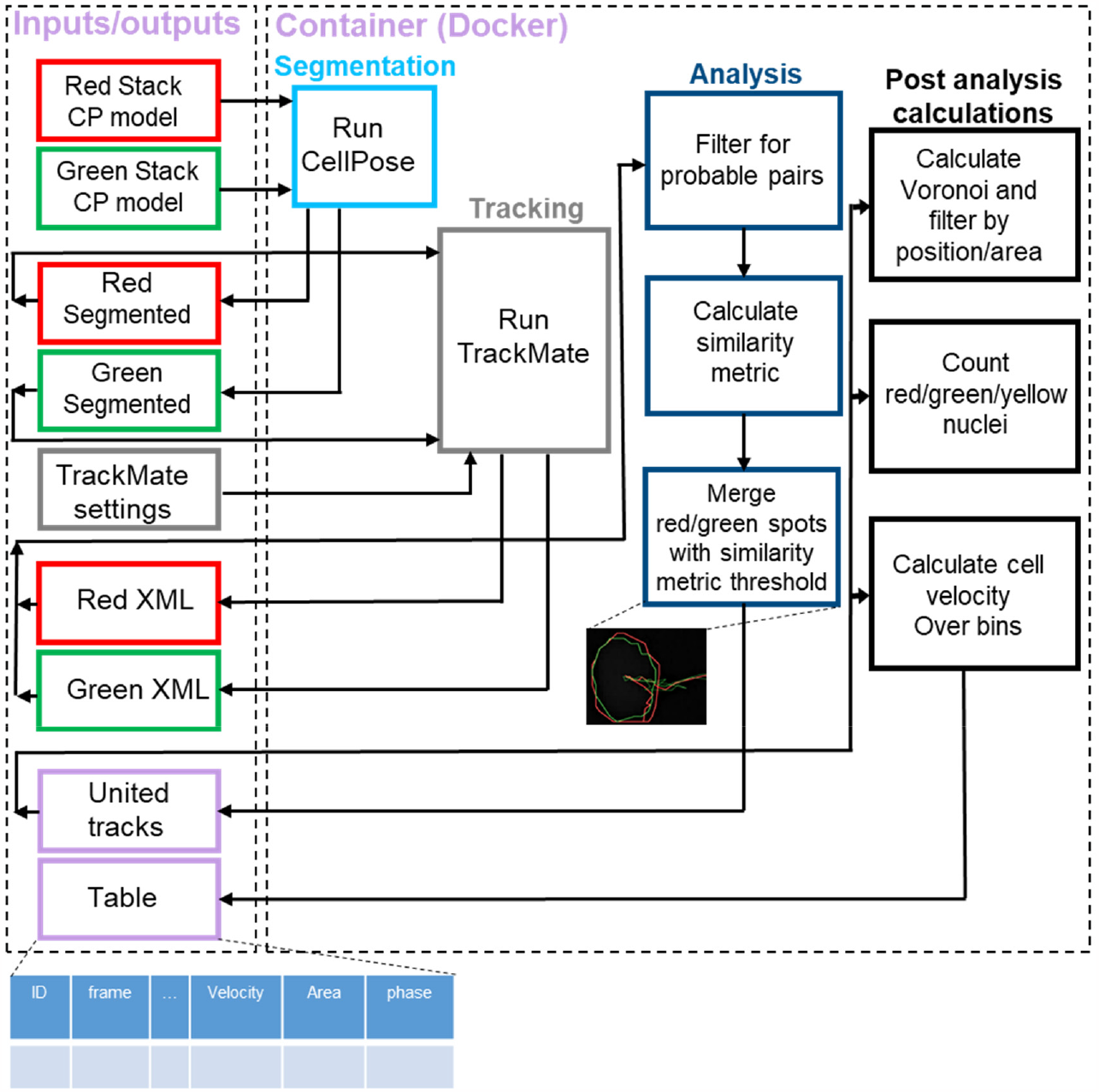
ConfluentFUCCI program flow chart. The flow chart is divided to two dashed rectangles. The left rectangle encircles the inputs and outputs of the program, when each stage output becomes the input for the next stage. The right rectangle encircles all applications, costume codes, and processes, used by the program and are wrapped in a container using Docker [14].

ConfluentFucci is available as open source in https://leogolds.github.io/ConfluentFUCCI/, but the quick and easier option is using the designed graphical user interface (GUI) (Fig. 3). The GUI is composed from the initiation screen, in which data is uploaded and initial settings are set, and the viewer screen which provides extremely easy and intuitive visualization of the segmented data. The installation and user guide are available in the supplementary material.

**Figure 3.**
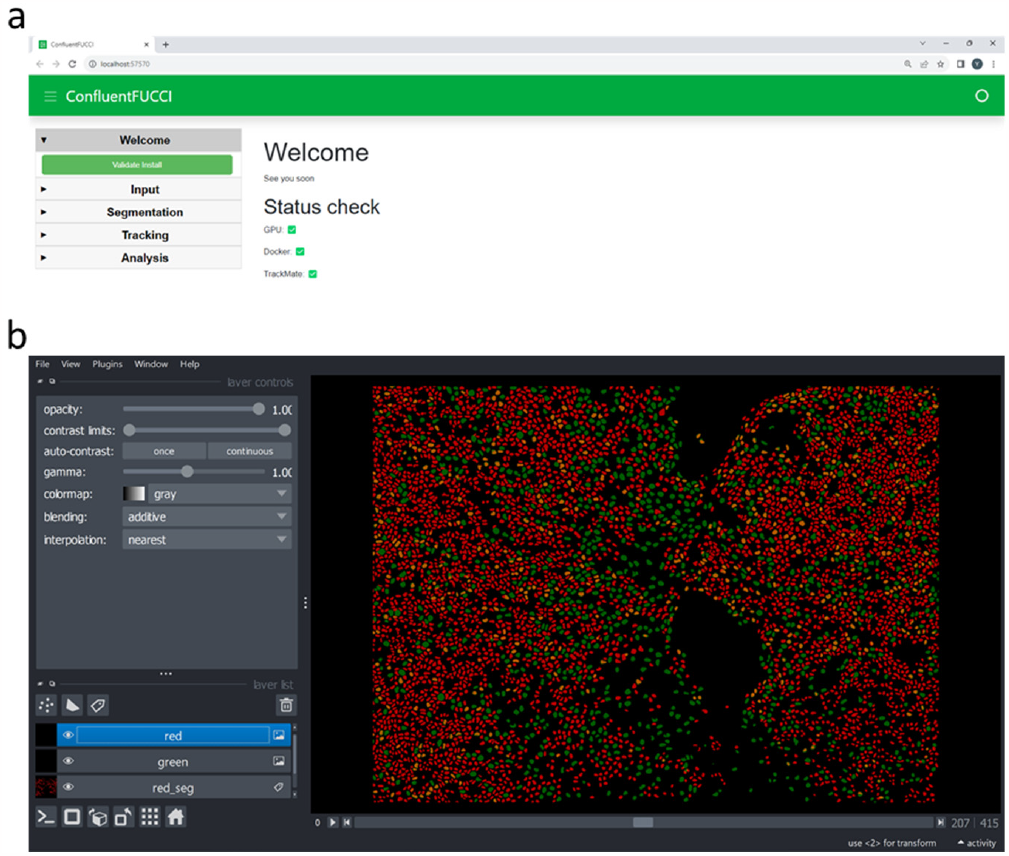
ConfluentFUCCI GUI. **a**, Initiation screen and, **b**, the viewer. The user guide, with exemplary data files to work with, are available in the supplementary material. See also supplementary video.

### Cell culture

MDCK II cells, transfected to stably express the FUCCI transgene, were a kind gift from Dr. Lars Hofnagel [4]. cells were cultured in Modified Eagle’s Medium (MEM, MERK), supplemented with 10% fetal bovine serum (FBS, MERK), 1% penicillin/streptomycin (PEN-STREP) and 1% L-glutamine (Sartorius). Cells were incubated at 37C, 85% humidity and 5% CO_2_. A 500 μm inner wall insert (Ibidi) was placed in the middle of a 35mm diameter TC petri dish. 56,000 cells were seeded in each well of the insert (2 in total). Cells were incubated for 24h, after which the insert was removed from the dish, to create a wound like gap between 2 confluent layers.

### Time-lapse fluorescent and PC microscopy

After insert removal, Petri dishes were secured under Zeiss AxioObserver 7 microscope with a stage top incubator (Ibidi), maintained at 37C, 85% humidity and 5% CO_2_. 10x, 3 channel images were obtained (phase contrast, green and orange), every 10 minutes. Green channel was obtained with the 38 HE filter set (470/525, Zeiss). Orange channel was obtained with the 43 HE filter set (550/605, Zeiss). To comply with FUCCItrack accepted input [8], the data used in Fig. 4a-c was acquired with Leica DMI 8 microscope, with the same culture conditions and a seeding density of 105,000 cells in each well.

**Figure 4.**
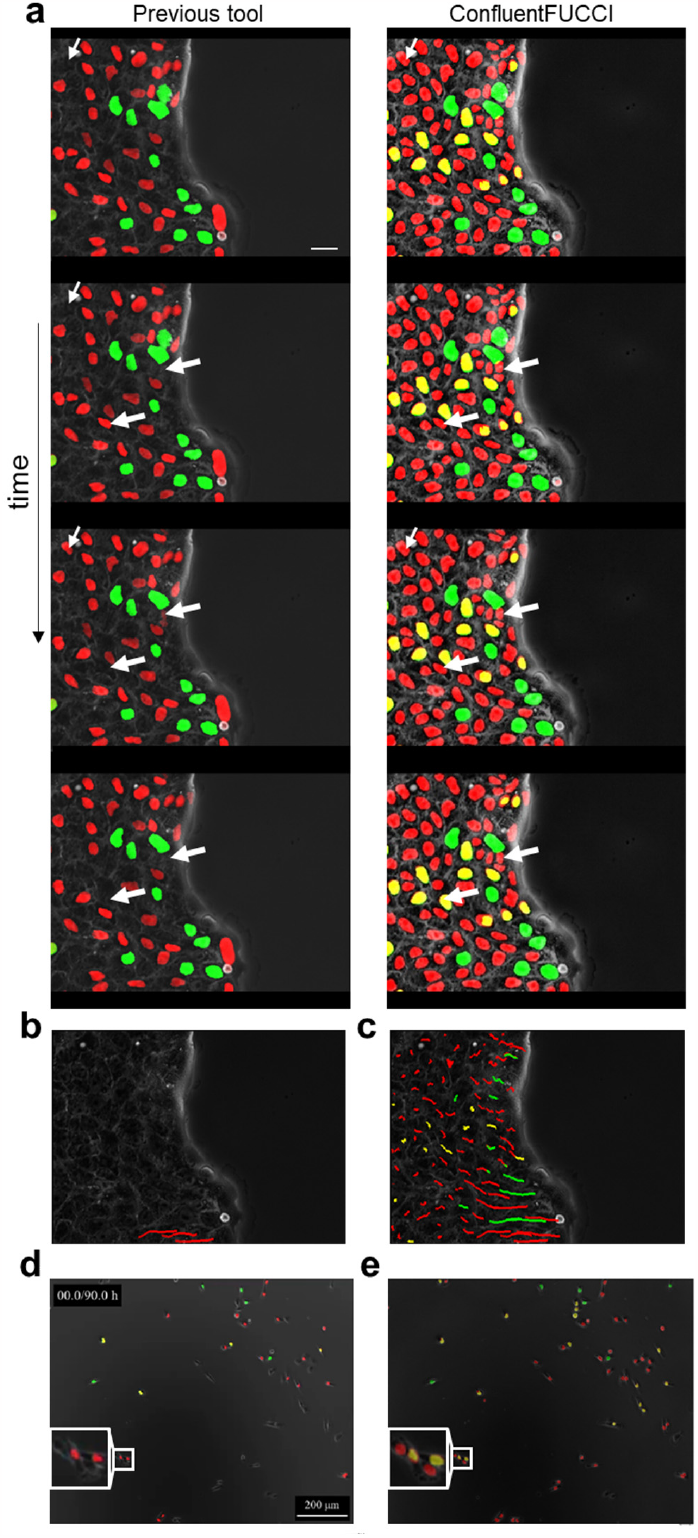
Robust segmentation and fully automatic time-tracking using ConfluentFUCCI. **a**, Confluent MDCK cells that collectively migrate in four sequential time points at 10 minutes intervals (see supplementary video). On the left, the segmentation done with the most recent published tool – FUCCItrack [8] which identifies only half of the population. On the right, the segmentation done with ConfluentFUCCI which identifies virtually all cells in the FOV. Thin white arrows indicate an instance in which a cell is identified at least 2 frames earlier in the right than in the left. Thick white arrows indicate a cell that is discontinuously identified on the left, while smoothly and continuously identified on the right. **b**, Three example of manually supervised tracking of cells (in **a)** using FUCCItrack, which took ∼5 min of per track. **c**, Automatically generated tracks of all cells (in **a)** using ConfluentFUCCI, which took ∼0.014 minutes of computation time per track. **d**, identification and segmentation of FUCCI cells in sparse culture of MDA-MB-231 done using FUCCItrack [8] results in many missed cells, as opposed to the results obtained using ConfluentFUCCI. Blowup indicates misidentification of FUCCI markers in regions where cell density increases.

**Figure 5.**
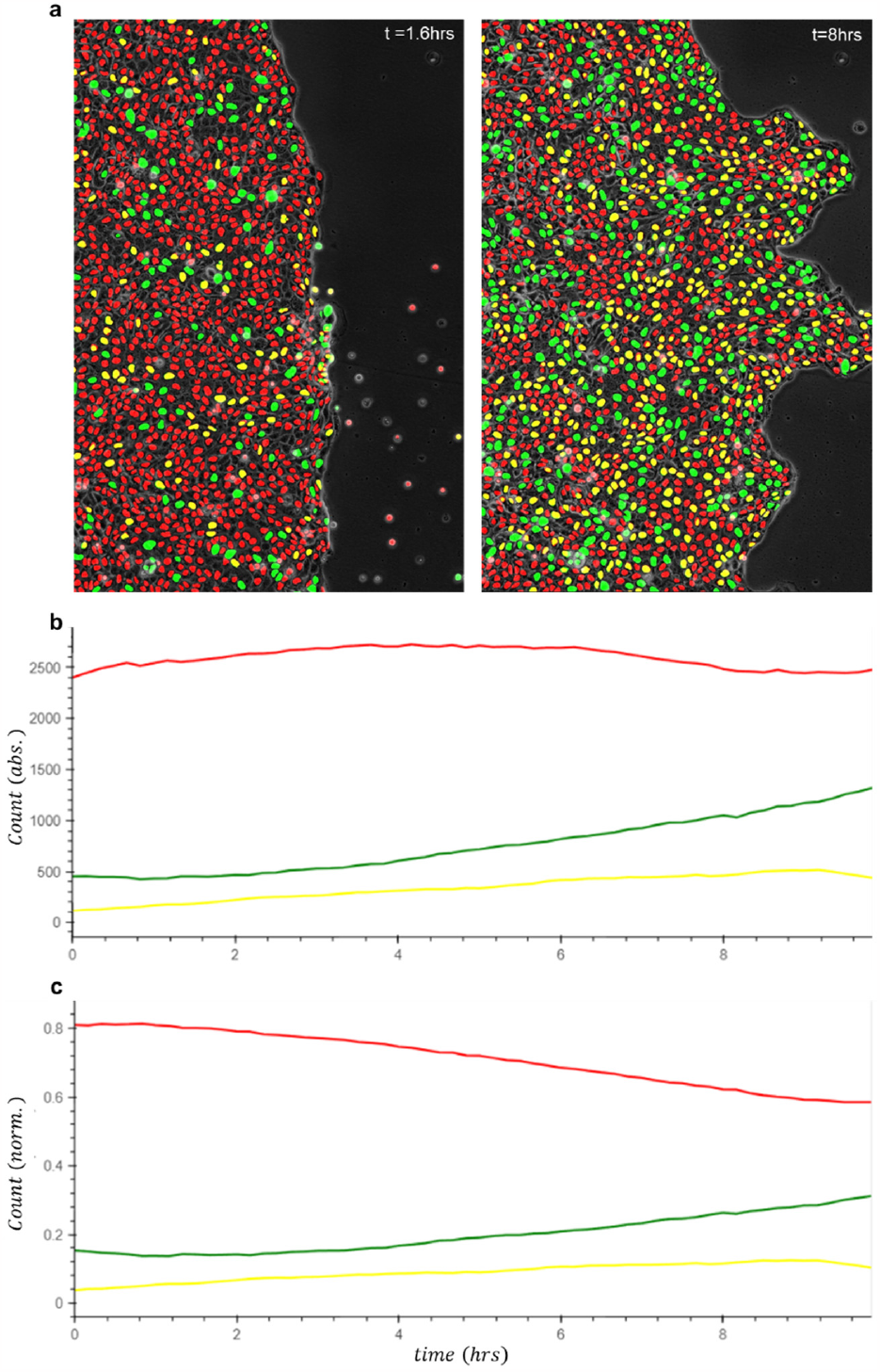
Tissue level monitoring of cell cycle progression during collective migration. **a**, Segmentation in two time points out of an analyzed stack of confluent MDCK cells that collectively migrate, post barrier lifting, for a 10 hours period. On the left t=1.6hrs, and on the right, t=10hrs. **b**, absolute number, and **c**, normalized number (fraction), of red\green\yellow tagged cells at each time point.

## Results

### Robust and fully-automated segmentation and time-tracking during cellular collective migration

As mentioned above, several tools have been uniquely developed for image analysis of FUCCI time-lapse datasets, with the most recent and advanced one being FUCCItrack [8, 10]. Although this tool does make some progress in bridging the existent literature gap for FUCCI data analysis, it suffers from a few substantial drawbacks in the context of analyzing a dense tissue that collectively migrate.

In FUCCItrack the segmentation process is based on the common, and above summarized, approach of combining adaptive thresholding, area-based filtering, and the morphological closing operation. We learned that this approach is indeed unsuitable for accurately segmenting and tracking FUCCI data in a dense tissue, after comparing the analysis performed by FUCCItrack vs. ConfluentFUCCI of our data of epithelial MDCK-FUCCI cells that collectively migrate. In a typical example shown in Fig. 4 (raw data available, and show in supplementary movie), we observe three major differences between the two analyses. First, Fig. 4a shows that within a single FOV the vast majority of cells are identified by ConfluentFUCCI as oppose to a substantial smaller fraction that is identified by FUCCItrack. Second, ConfluentFUCCI continuously detect all nuclei throughout the entire duration of the experiment (Fig. 4a), whereas, FUCCItrack discontinuously detect many nuclei that flicker on and off (Fig. 4a; white arrows). Third, all time tracks for nuclei positions are automatically produced by ConfluentFUCCI (Fig. 4c). However, FUCCItrack requires the user to manually choose each single nucleus to be detected, and furthermore to supervise the tracking process frame by frame (Fig. 4b). We manually tracked multiple cells (in Fig. 4d) and measured ∼5 minutes per track of labor time, which scale to a staggering time of ∼8 hours for just this one position of highly sparse condition, in just one experiment. For comparison, we analyzed our data with ∼11k cells, and the entire analysis took 2.5 hrs, which means ∼0.014 minutes per track. We made further comparison for even sparse systems using available data previously analyzed with FUCCItrack [8]. Fig. 4d shows a snapshot of a sparse culture of MDA-MB-231 cell line that is stably transfected for the FUCCI transgene. Interestingly, ConfluentFUCCI was able to detect all nuclei in the image, with FUCCItrack identifying less than half, and misidentification seemed to occur in high cell density locations (Fig. 4d; blowup).

### Tissue-level monitoring for cell cycle status during collective migration

The most straightforward use of the FUCCI system is to asses viability and global proliferation patterns at the tissue level [6, 24]. For example, monitoring the ratio of cells that transition to S state in one culture sample that was pharmacologically manipulated, or implanted and then extracted from in situ growth [25]. The FUCCI system can then provide a powerful method to evaluate cell cycle related mechanisms, with the most pronounced one being studies of cell cycle targeted drugs for cancer growth suppression and metastatic inhibition [26-28]. Famous metastatic hallmarks are the epithelial to mesenchymal transition (EMT), or partial EMT, and resultant cellular migration capacity [29-31]. Hence, even without considering single-cell level analysis, just a robust computerized tool that counts, per FOV, the number of cells in total, and number of cycling cells, may provide great clinical contribution. Fig. 5a&b demonstrate the ability of ConfluentFUCCI to accurately segment nuclei, and identify cell cycle status, in virtually all cells in a collective migration experiment of MDCK-FUCCI cells throughout a 10hrs duration. Not like previous studies which intentionally design the culture for extremely low density and counts about a hundred cells per FOV [8, 10], in Fig. 5c we show how the ConfluentFUCCI counts about three thousand cells per FOV. This gives a remarkable statistical power, and the ability to clearly define the global proliferation trend in the migrating tissue. Consistent with previous studies [4, 32], at early times the number of cells in total, and the number of cycling cells do not change (Fig. 5c,d). This shows that the initial expansion of the tissue is dominated solely by collective cellular migration and not by proliferation. After about two hours of collective migration the fraction of non-cycling cells start decreasing, with a directly compatible increase of the fraction of cycling cells (Fig. 5d). It is important to note that this type of extremely accurate (Fig .5) cell cycle analysis at the tissue-level, is of course not limited to dynamic processes and can be used for fixed samples or histological sections with high cellular density.

### Single-cell-level monitoring for cell cycle status, dynamics, and morphology

The regulation of cell cycle progression by either intracellular biochemical cues, or intercellular mechanical cues, has been extensively studied for many years now [4, 5, 33, 34]. Nevertheless, when it comes to cell cycle regulation during tissue expansion and cellular collective migration, it is still unclear what are the main contributing molecular or mechanical factors. Many key mechanical factors, such as intercellular stress, cell morphology, or growth rate, are inherently related to sterical constraints imposed on each cell by its multicellular neighborhood. Unlike current tools [8, 10], ConfluentFUCCI allows for a robust and fully automated monitoring of all nuclei during collective migration and enable investigating the spatio-temporal connection between cell cycle progression and a variety of participating mechanical factors. To demonstrate such investigation, we decided to explore the relationship between cell area and the transition from G1 to S state in collectively migrating epithelial cells. Fig. 6a&b shows how the condition of a condensed cellular environment allows for a good size estimation of each cell using Voronoi tessellation that is based on the nuclei positions.

**Figure 6.**
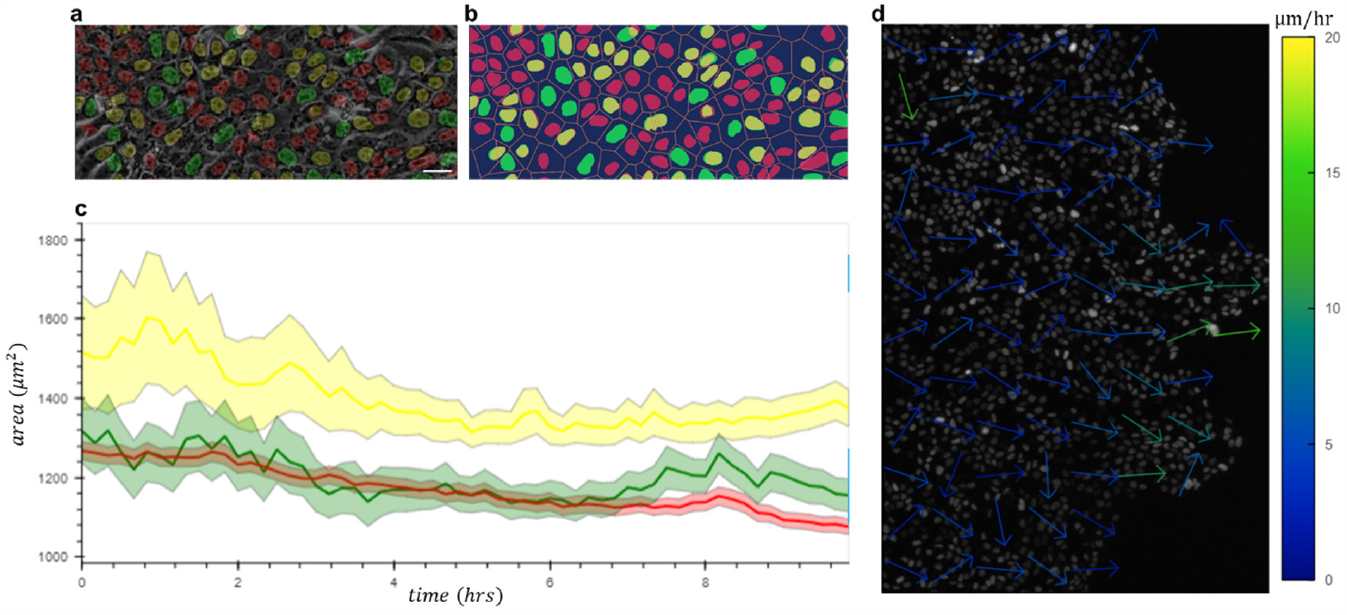
Single cell level tracking of all cells within an advancing tissue, that is confluent and dense, allows measuring cellular morphology and velocity field. **a**, Confluent MDCK cells that collectively migrate, post barrier lifting (as in Figs. 4a&5), with fluorescent FUCCI tags overlaid on the phase contrast (PC) image. Scale bar, 50 μm. **b**, The area of each cell is approximated using Voronoi tessellation that is based on nuclei centers. **c**, Solid lines, colored according to cell cycle state, show the average area of cells in each state. Colored shaded regions indicate 95% confidence intervals. **d**, Velocity vector field based on nuclear velocity.

During the entire expansion process of the tissue we used ConfluentFUCCI to track all nuclei and define which emitted, only on the MKO2 channel (G1 state), only on the mAG1 channel (S/G2/m state), or in both MKO2 and mAG1 (G1 to S transition). Fig. 6d shows that the area of cells that are transitioning from G1 to S are systematically bigger. This essentially replicates previously published [4]data, according to which the authors implied that a mechanosensitive checkpoint ensures that only cells big enough, and that can physically facilitate all mitosis related events will progress in the cell cycle. To establish the existence of that area related checkpoint in greater depth one needs to further examine the correlation between the transition event to other, molecular, or physical, temporal features of the transitioning cells. Many of this physical features can be investigated by ConfluentFUCCI and include, for example, all morphological characteristics of each cell and its nucleus, their trajectories, and the Eulerian-like velocity field in the layer. Fig 6c shows the velocity field, which is possible to calculate only because ConfluentFUCCI tracks all cells in this high cellular density conditions, and in essence transform Lagrangian trajectories to an Eulerian velocity field.

## Discussion

This work presents a new fully-automated image analysis tool for analyzing live fluorescent markers for cell cycle monitoring (FUCCI), in time-lapse microscopy data of cells that collectively migrate on flat surface. The tool is capable of accurately segmenting and tracking nuclei in dense tissues and identifying the cell cycle stage of individual cells. In comparing ConfluentFUCCI with the most recent and relevant existing tool, we found that ConfluentFUCCI outperforms it in terms of segmentation ability, accuracy, and time tracking, both in sparse and dense environments. Our work also demonstrates the ability of ConfluentFUCCI to monitor cell cycle status and dynamics at both the tissue and single-cell levels, as well as replicate the main features of the previously published relationship between cell cycle progression and cell size. We suggest that ConfluentFUCCI can provide valuable, reproducible and user independent insights into cell cycle regulation in dense cellular condition in the context of various biological disciplines including cancer growth and metastasis. We would like to note that ConfluentFUCCI was developed as part of a current effort we take to integrate, on a large scale of data, measurements of cell cycle dynamics, ECM-cell and cell-cell mechanical stresses, in different crowded cancerous environments.

## Availability Statement

Data, code, and software are available on github repository, https://leogolds.github.io/ConfluentFUCCI/. The available data is an integral part of the installation process and is immediately available for the user after installation. The data is also available for direct download at https://github.com/leogolds/ConfluentFUCCI/tree/main.

## Acknowledgments

This work was supported by: the Pearlstone Center at Ben-Gurion University; the Israeli Science Foundation (ISF) on the individual research grant number 2107/21, and the New-Faculty Equipment grant number 2108/21.

## References

1. Wiman, K.G. and B. Zhivotovsky, Understanding cell cycle and cell death regulation provides novel weapons against human diseases. Journal of Internal Medicine, 2017. 281(5): p. 483–495.

2. Benham-Pyle, B.W., B.L. Pruitt, and W.J. Nelson, Mechanical strain induces E-cadherin-dependent Yap1 and beta-catenin activation to drive cell cycle entry. Science, 2015. 348(6238): p. 1024–1027.

3. Rodríguez-Franco, P., et al., Long-lived force patterns and deformation waves at repulsive epithelial boundaries. Nature Materials, 2017. 16: p. 1029.

4. Streichan, S.J., et al., Spatial constraints control cell proliferation in tissues. Proceedings of the National Academy of Sciences, 2014. 111(15): p. 5586–5591.

5. Uroz, M., et al., Regulation of cell cycle progression by cell–cell and cell–matrix forces. Nature Cell Biology, 2018. 20(6): p. 646–654.

6. Zielke, N. and B.A. Edgar, FUCCI sensors: powerful new tools for analysis of cell proliferation. Wiley Interdisciplinary Reviews: Developmental Biology, 2015. 4(5): p. 469–487.

7. Sakaue-Sawano, A., et al., Visualizing Spatiotemporal Dynamics of Multicellular Cell-Cycle Progression. Cell, 2008. 132(3): p. 487–498.

8. Taïeb, H.M., et al., FUCCItrack: An all-in-one software for single cell tracking and cell cycle analysis. PLOS ONE, 2022. 17(7): p. e0268297.

9. Roccio, M., et al., Predicting stem cell fate changes by differential cell cycle progression patterns. Development, 2013. 140(2): p. 459–470.

10. Ghannoum, S., et al., CellMAPtracer: A User-Friendly Tracking Tool for Long-Term Migratory and Proliferating Cells Associated with FUCCI Systems. Cells, 2021. 10(2): p. 469.

11. Wen, T., et al., Review of research on the instance segmentation of cell images. Computer Methods and Programs in Biomedicine, 2022. 227: p. 107211.

12. Minaee, S., et al., Image Segmentation Using Deep Learning: A Survey. IEEE Transactions on Pattern Analysis and Machine Intelligence, 2022. 44(7): p. 3523–3542.

13. Stringer, C., et al., Cellpose: a generalist algorithm for cellular segmentation. Nature Methods, 2021. 18(1): p. 100–106.

14. Ershov, D., et al., TrackMate 7: integrating state-of-the-art segmentation algorithms into tracking pipelines. Nature Methods, 2022. 19(7): p. 829–832.

15. Merkel, D., Docker: lightweight Linux containers for consistent development and deployment. Linux J., 2014. 2014(239): p. Article 2.

16. Harris, C.R., et al., Array programming with NumPy. Nature, 2020. 585(7825): p. 357–362.

17. eam, T.p.d., pandas-dev/pandas: Pandas. Zenodo, 2020.

18. Rocklin, M., Dask: Parallel Computation with Blocked algorithms and Task Scheduling. Proceedings of the 14th Python in Science Conference, 2015.

19. Samuels; stonebig; Florian LB; Andrew Tolmie; Daniel Stephan; Scott Lowe; John Bampton; henriqueribeiro; Irv Lustig; Julia Signell; Justin Bois; Leopold Talirz; Lukas Barth; Maxime Liquet; Ram Rachum; Yuval Langer; arabidopsis; kbowen, P.R.J.-L.S.J.A.B.B.N.A.C.B.A.R.J.M.V.T.m.M.K.e.g.J., holoviz/holoviews: Version 1.13.3. Zenodo, 2020.

20. team, P.d., Panel: The powerful data exploration & web app framework for Python. https://panel.holoviz.org/.

21. Chiu, C.-L. and N. Clack, napari: a Python Multi-Dimensional Image Viewer Platform for the Research Community. Microscopy and Microanalysis, 2022. 28(S1): p. 1576–1577.

22. Kaliman, S., et al., Limits of Applicability of the Voronoi Tessellation Determined by Centers of Cell Nuclei to Epithelium Morphology. Frontiers in Physiology, 2016. 7(551).

23. Saraswathibhatla, A., et al., Coordinated tractions increase the size of a collectively moving pack in a cell monolayer. Extreme Mechanics Letters, 2021. 48: p. 101438.

24. Yano, S., et al., FUCCI Real-Time Cell-Cycle Imaging as a Guide for Designing Improved Cancer Therapy: A Review of Innovative Strategies to Target Quiescent Chemo-Resistant Cancer Cells. Cancers, 2020. 12(9): p. 2655.

25. Lu, Y., et al., Evaluating the Accuracy of FUCCI Cell Cycle In Vivo Fluorescent Imaging to Assess Tumor Proliferation in Preclinical Oncology Models. Molecular Imaging and Biology, 2022. 24(6): p. 898–908.

26. Sherr, C.J. and J. Bartek, Cell Cycle–Targeted Cancer Therapies. Annual Review of Cancer Biology, 2017. 1(1): p. 41–57.

27. Schwartz, G.K. and M.A. Shah, Targeting the Cell Cycle: A New Approach to Cancer Therapy. Journal of Clinical Oncology, 2005. 23(36): p. 9408–9421.

28. Diaz-Moralli, S., et al., Targeting cell cycle regulation in cancer therapy. Pharmacology & Therapeutics, 2013. 138(2): p. 255–271.

29. Wong, I.Y., et al., Collective and individual migration following the epithelial– mesenchymal transition. Nat Mater, 2014. 13(11): p. 1063–1071.

30. Mitchel, J.A., et al., In primary airway epithelial cells, the unjamming transition is distinct from the epithelial-to-mesenchymal transition. Nature Communications, 2020. 11(1): p. 5053.

31. Hanahan, D. and Robert A. Weinberg, Hallmarks of Cancer: The Next Generation. Cell, 2011. 144(5): p. 646–674.

32. Trepat, X., et al., Physical forces during collective cell migration. Nature Physics, 2009. 5(6): p. 426–430.

33. Watt, F.M., P.W. Jordan, and C.H. O’Neill, Cell shape controls terminal differentiation of human epidermal keratinocytes. Proceedings of the National Academy of Sciences, 1988. 85(15): p. 5576–5580.

34. Folkman, J. and A. Moscona, Role of cell shape in growth control. Nature, 1978. 273(5661): p. 345–349.

